## Supplementary figures and images for "Human adherent cortical organoids in a multiwell format"

### supplementary figure 1

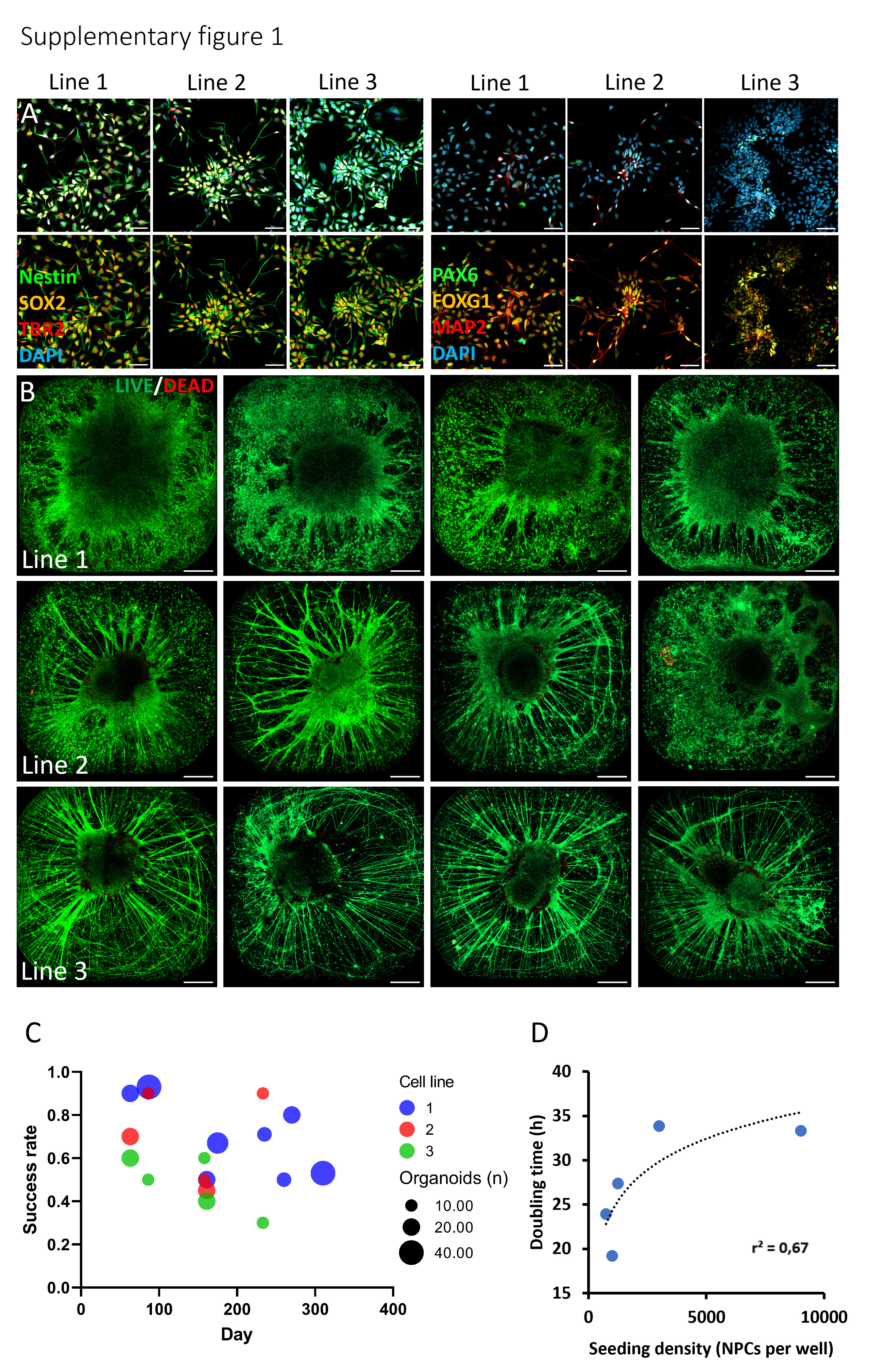

### supplementary figure 2

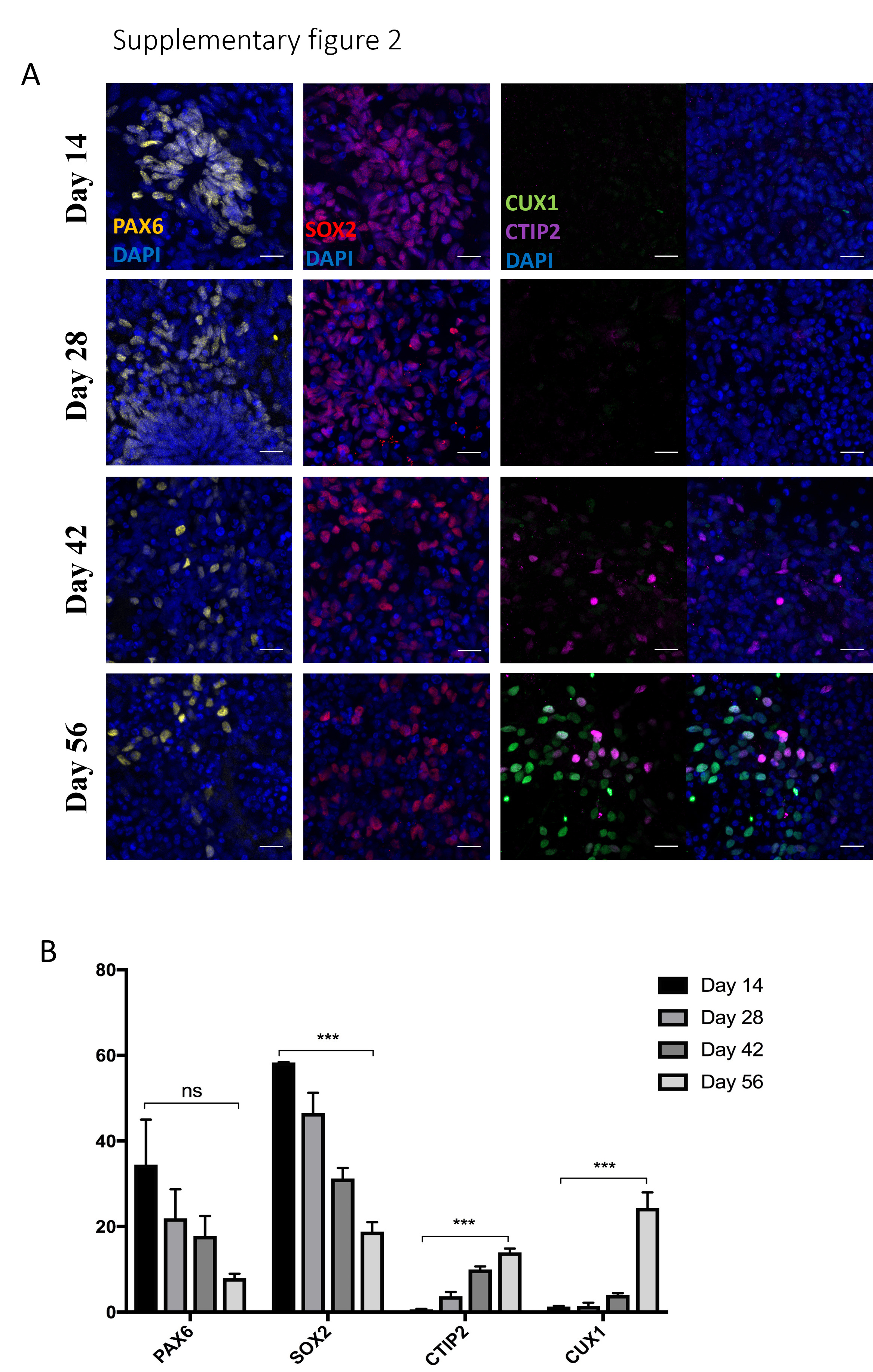

### supplementary figure 3

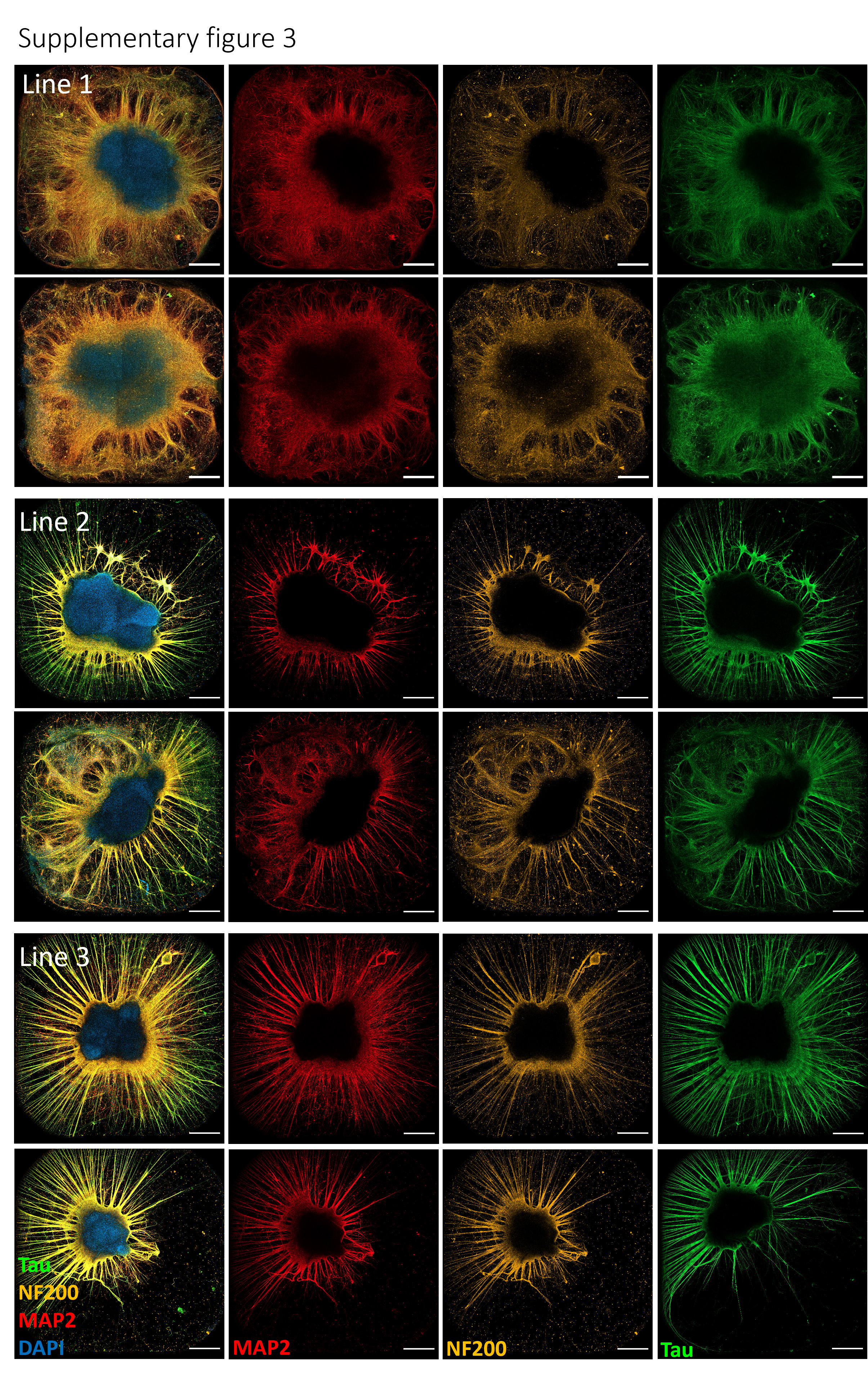

### supplementary figure 4

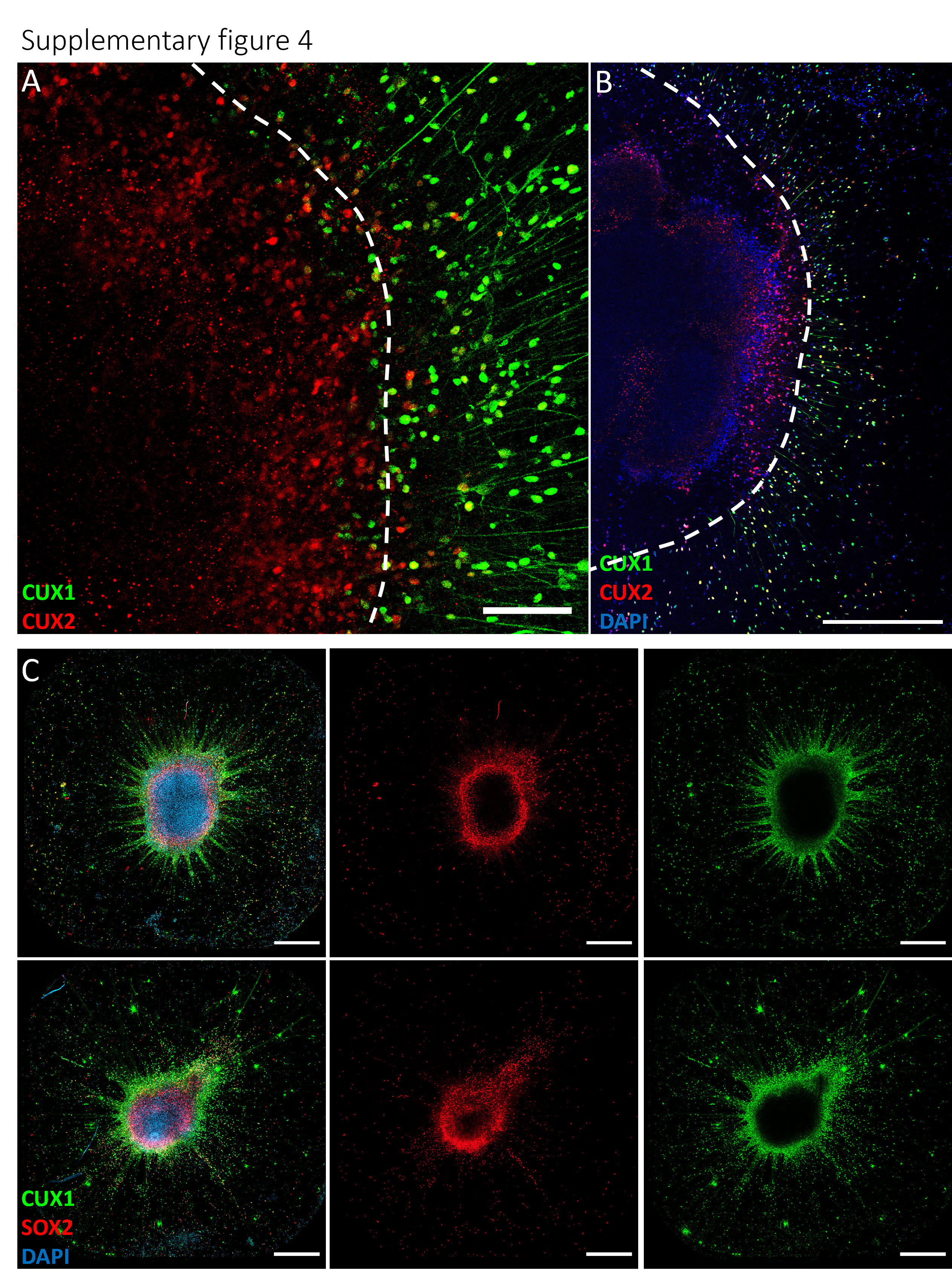

### supplementary figure 5

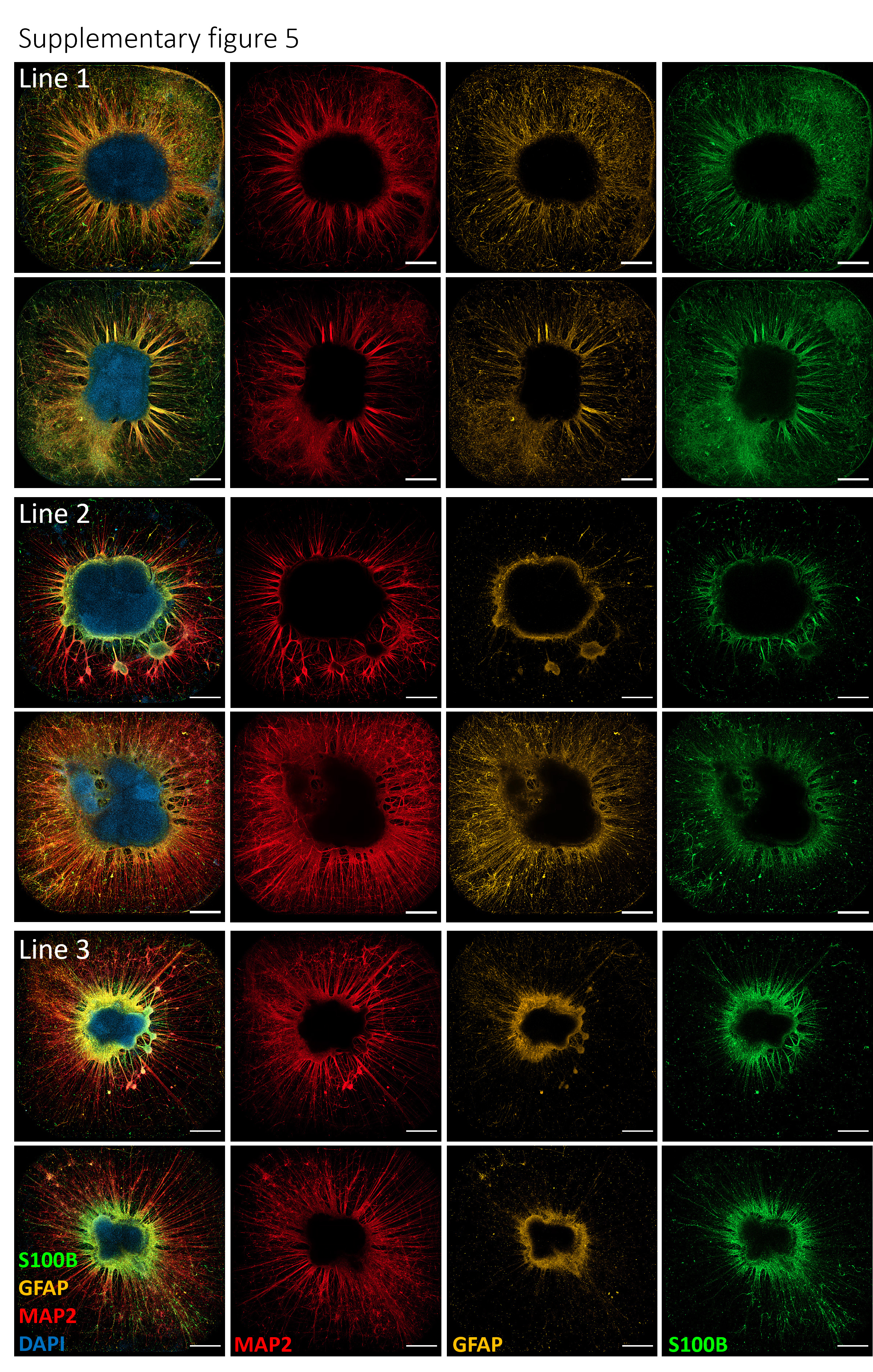
